# Dynamic Compression Platform for Live Imaging of Scaffold-Transmitted Cellular Mechanoresponses

**DOI:** 10.64898/2026.09.09.749723

**Authors:** Marie Moulin, Elisa G. Bissacco, Stephen J. Ferguson, Paul O’Callaghan

## Abstract

Mechanical characterization of biomaterial scaffolds is essential to evaluate their capacity to meet the functional demands of target tissues in tissue engineering and regenerative medicine applications. Scaffolds designed to interface with living tissues must support the transmission of mechanical cues to resident cells and stimulate mechanosignaling pathways that are essential to their function. In joints, bone and cartilage cells act as primary mechanosensors, converting mechanical stimuli into biochemical signals that regulate tissue homeostasis and remodelling. Therefore, evaluating cellular mechanoresponses to scaffold-transmitted compression *in vitro* can inform the development of functional tissue-engineered constructs. For example, poly(ε-caprolactone) (PCL) scaffolds are highly relevant for bone and cartilage tissue engineering due to their biocompatibility, stable mechanical properties and slow degradation. Here, we applied a custom-built device to study compression-induced mechanosignaling in MC3T3-E1 pre-osteoblast cells. The device is composed of a polydimethylsiloxane (PDMS) pillar, a force-sensing load cell, and a piezoelectric linear track. A protocol is described in which MC3T3-E1 cells are repeatedly compressed, while in parallel live tracking of force measurements and live imaging of intracellular calcium dynamics in MC3T3-E1 cells are recorded. PCL scaffolds fabricated by melt electrowriting (MEW) were subsequently integrated into the platform. Scaffold-transmitted compression triggered dynamic increases in cytosolic Ca²⁺ in MC3T3-E1 cells located directly under the PCL microfibers, but also in cells located in the interfiber spaces. This device and workflow facilitate *in vitro* investigations of real-time cellular mechanoresponses to dynamic compression applied with biomaterial scaffolds, and provides a testing platform for evaluating the mechanotransductive properties of scaffolds intended for tissue engineering applications.

## Introduction

Abnormal loading conditions arising from factors such as immobilization, obesity, or trauma can alter joint tissue composition, structure, metabolism, and mechanical properties, ultimately contributing to pathological conditions such as osteoarthritis^1^. Tissue engineering strategies based on biomaterial scaffolds aim to generate physiologically relevant tissue substitutes for the clinical repair and replacement of damaged joint tissues^2^. Various fabrication techniques have been developed to produce three-dimensional (3D) scaffolds designed to mimic the structural and functional properties of the extracellular matrix (ECM). Among these approaches, melt electrowriting (MEW) is an additive manufacturing technique that enables the fabrication of micro-scale fibrous scaffolds with precisely controlled architecture using biocompatible polymers such as poly(ε-caprolactone) (PCL)^3,4^. MEW-fabricated PCL scaffolds have excellent biocompatibility, mechanical stability, and slow biodegradation rates, and have been investigated for their potential applications in diverse tissues, including cartilage and bone^5,6^. While the mechanical properties of scaffolds are commonly characterized to determine their relevance for load-bearing tissue applications, less is understood about how cells interacting with these scaffolds respond to mechanical loading *in vivo*. This highlights the need for *in vitro* testing platforms capable of investigating cellular responses to scaffold-transmitted mechanical stimulations.

Cells sense mechanical stimuli through various mechanisms, including mechanosensitive ion channels (MICs), which convert physical forces, such as changes in membrane tension, into biochemical signals^7^. These mechanical cues act as key biophysical regulators of cell fate, including proliferation, differentiation, and apoptosis, thereby maintaining tissue homeostasis through pathways such as YAP/TAZ^8,9^. MICs such as Piezo1 and Transient Receptor Potential Vanilloid 4 (TRPV4) primarily gate the influx of Ca²⁺ ions from the extracellular space to the cell cytosol, triggering downstream signaling events and gene regulation^10–14^. In osteoblastic cells, calcium signaling is a critical regulator of processes such as osteogenic differentiation^15^, and studies have identified Piezo1 and TRPV4 as key mechanotransducers essential for bone adaptation to loading^16–18^. Therefore, investigating cellular calcium responses provides insight into transmission of mechanical cues from scaffolds to cells, particularly as appropriate mechanical stimulation is required to regulate osteoblastic cell behavior and maintain tissue integrity^19,20^. *In vitro*, mechanoresponses in cells can be investigated using different techniques, all of which induce changes in plasma membrane tension, including atomic force microscopy (AFM), pillar arrays, glass probes, microfluidic devices, and ultrasound^21–23^.

In this study, a custom uniaxial compression tool, previously applied to compress tumor cells and isolated pancreatic islets^24,25^, was used to investigate calcium responses of pre-osteoblastic cells subjected to compression, in contact with PCL scaffolds. The compression device consists of a polydimethylsiloxane (PDMS) pillar fitted in a donut-shaped force-sensing load cell, which is connected and actuated via a linear piezoelectric track, and mounted on a confocal microscope stage. The position of the PDMS compression pillar can be controlled vertically with submicron precision via the linear piezoelectric track, and the force exerted on the cell layer can be measured via the donut load cell^24^. The imaging of cytosolic calcium dynamics in response to compression was recorded with live confocal imaging using established fluorescent calcium reporters. MC3T3-E1 pre-osteoblastic cells exhibited transient calcium responses upon mechanical stimulation via PDMS pillar compression. We confirmed that the MC3T3-E1 cultures were sensitive to established pharmacological agonists of Piezo1 and TRPV4 mechanosensitive ion channels, suggesting that these were among the mechanosensors eliciting cell responses to pillar-mediated compression. The compression device was then applied to study the mechanotransductive properties of melt-electrowritten PCL scaffolds, when used as the contacting countersurface to the cells, which present relevance for potential application in bone and cartilage tissue engineering^26^. The optical clarity of the PDMS pillar permitted visualization of the PCL scaffold during cell compression, while parallel confocal live-imaging of fluorescence Ca²⁺ reporters recorded individual cell responses. MC3T3-E1 cells were subjected to repeated compression events, and increases in cytosolic Ca²⁺ were detected for cells directly compressed under the PCL microfibers and for cells located within the interfiber spaces. Collectively, these findings establish the platform as a versatile approach for interrogating real-time cellular mechanoresponses to biomaterial scaffolds under dynamic compression protocols. More broadly, this workflow provides a novel framework for evaluating how scaffold architecture and mechanical properties influence mechanotransduction with relevance to tissue engineering applications.

## Results and Discussion

### Characterization of the Cell Press tool for controlled compression

The *in vitro* mechanostimulation platform used in this study is based around a previously developed cell compression device^24^, and referred to as the Cell Press. It integrates a linear piezoelectric track, a donut load cell, and a polydimethylsiloxane (PDMS) compression pillar (Fig. 1A). Custom-designed 3D-printed components enable assembly of the system and mounting onto the microscope stage rail (Fig. 1B), ensuring precise alignment of the PDMS pillar within a cell culture dish (Fig. 1C). The PDMS pillar is cast from a custom 3D-printed mold and actuated along its vertical axis via the piezoelectric track, allowing direct mechanical compression of cells. In parallel, the donut-shaped load cell allows real-time measurement of the applied compression forces. Integration of the device on a confocal microscope stage enables synchronized mechanical stimulation and live imaging, and the transparency and biocompatibility of the PDMS pillar make it well suited for imaging applications^23^.

**Figure 1.**
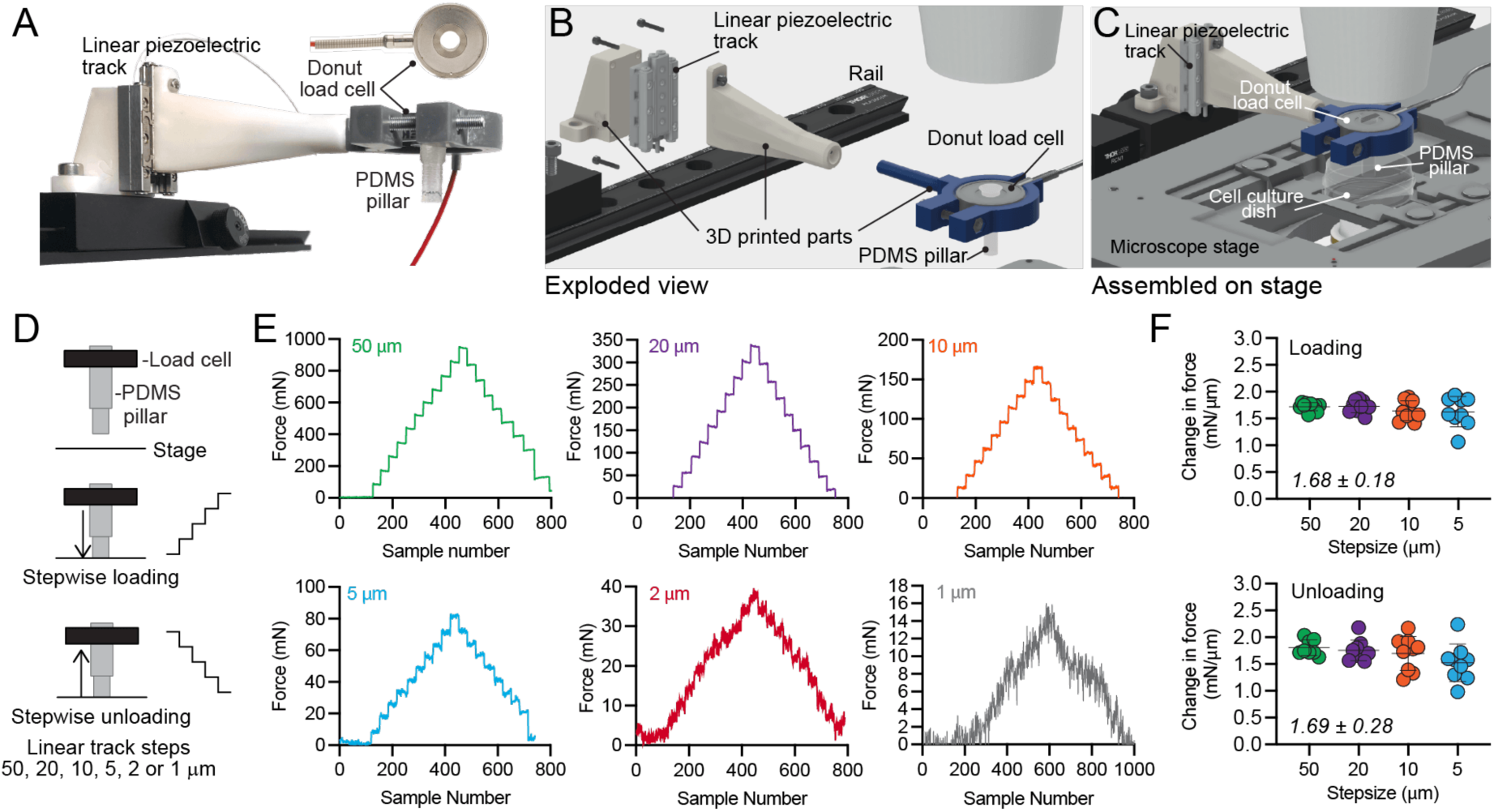
Overview of the assembly and operation of the cell compression tool. **A.** Photograph of the assembled cell compression press tool, previously introduced in^24^. The position of the linear piezoelectric track, the donut-shaped load-sensing cell, and the PDMS compression pillar are illustrated. **B.** Exploded view of the CAD drawing of the cell compression tool, indicating the custom designed and 3D-printed parts. **C.** CAD drawing of the full assembly of the cell compression tool on the confocal microscope stage with a cell culture dish in place. **D.** Overview of the protocol applied to load and unload the PDMS pillar directly against the microscope stage. The tool was assembled as in A, and the controller for the linear piezoelectric track was pre-set for each of the indicated step sizes. The controller was manually operated to displace the pillar downward during loading and upward during unloading in a stepwise manner. **E.** Dynamic force measurements were acquired at a rate of 1 sample/second with the Futek donut load cell interfaced with a USB signal conditioner, with signals visualized and logged using Sensit data acquisition software. Each panel represent the recorded forces (mN) during PDMS pillar loading and unloading for each of the indicated step sizes (50 μm – 1 μm). **F.** Quantification of the average change in force (mN)/μm, calculated for each of the indicated step sizes (50 μm – 5 μm) during the loading phase (upper plots) and unloading phase (lower plots). One-way ANOVA with multiple comparisons detected no significant differences between the average change in force/μm between the step sizes. The average change in force (mN/μm) + standard deviation for all quantified step sizes is presented in the respective plots.

The compression performance of the Cell Press was evaluated using controlled loading and unloading protocols on the microscope stage with defined displacement step sizes ranging from 50 ∝m to 1 μm (Fig. 1D), which were set and manually adjusted via the controller operating the piezoelectric track. The PDMS pillar was displaced downward onto the solid microscope stage for stepwise loading and upwards for stepwise unloading via the piezoelectric controller, with the pre-set steps incrementally applied at regular time intervals (Fig. 1D). Force (mN) profiles registered throughout the loading and unloading cycles showed that displacements taken with each step size produced a measurable force, and distinct changes in the force measurements were clearly discernible down to 5 ∝m step-sizes, still evident with a 2 ∝m step size, but became difficult to distinguish with a 1 μm step size, although changes in force were still clearly detectable (Fig. 1E). This force response arises from elastic deformation of the PDMS pillar^27^, which accommodates the imposed axial compression exerted between the microscope stage and the inner surface of the donut load cell. Across displacement step sizes from 50 to 5 μm, force increased proportionally with displacement, resulting in consistent force generation per micrometer during loading (1.68 ± 0.18 mN/μm) and unloading (1.69 ± 0.28 mN/μm), with no significant differences between conditions (Figure 1F). Together, these results demonstrate that compression of the elastic PDMS pillar enables reproducible, and consistent force generation across a range of displacement step sizes.

When applied to cells in a culture dish, the Cell Press tool is intended to apply controlled mechanical stimulation to cells beneath the pillar during live imaging experiments. Several approaches have been developed to mechanically stimulate cells *in vitro* using controlled compressive loading while imaging responses^28^. These include atomic force microscopy (AFM) based methods and engineered microdevice-based systems^23^. AFM enables precise application of localized forces to single cells or spheroids via indentation while also allowing measurement of cell mechanical responses^21,29^. Microdevices instead integrate actuators to control the compression of cells cultured as monolayers or in three-dimensional cultures^23,30,31^. Among these microdevices, the versatility of microfluidic platform design and their compatibility with on-chip microscopy have facilitated diverse strategies, including pneumatic actuation of deformable PDMS diaphragms^32,33^ or PDMS-based micro-piston systems^31,34^. The Cell Press represents an easily adaptable complement to existing mechanostimulation approaches by integrating controlled compression, direct force measurement, and live imaging compatibility into a compact and customizable platform, and previously proven to be compatible with compression of monolayers and organoid-sized microtissues such as islets^24,25^.

### MC3T3-E1 cell responses to mechanical compression and mechanosensitive ion channel agonists

The Cell Press was applied to compress pre-osteoblastic MC3T3-E1 cells via direct repeated contact with the surface of the PDMS pillar, while simultaneously monitoring cells expressing a fluorescent calcium reporter (Fig. 2A and B). To mechanically induce Ca^2+^ responses, the pillar was displaced downward towards the cell layer with 2 µm step increments. Initial contact with the cell layer was confirmed via the live force measurement recordings; however, additional force was required to induce an increase in cytosolic Ca^2+^ (Fig. 2C-D) in the imaged cell. Several compression cycles were applied, resulting in rapid and transient increases in cytosolic Ca^2+^, with each peak response occurring at approximately the same compression force (40 mN), which was achieved via multiple 2 µm compression steps. The kymograph presented in Fig. 2E illustrates the spatiotemporal dynamics of the Ca²⁺ signal along a defined axis within the cell (pink arrow, Fig. 2B), revealing the rapid increase, propagation and duration of Ca^2+^ responses during the successive compression cycles (Fig. 2E). Despite sustained compression following a Ca^2+^ response, the signal progressively declined toward baseline. The transience of the Ca^2+^ increase may reflect an accommodation response, whereby cells adapt their morphology and associated membrane tension during sustained compression^35,36^. Alternatively, this may reflect the presence of desensitization mechanisms known to inactivate mechanosensitive ion channels even in the presence of sustained stimulus^37^. Nonetheless, once the pillar was retracted, it was possible to repeatedly elicit compression-induced Ca^2+^ responses at similar loads (Fig. 2C, D).

**Figure 2.**
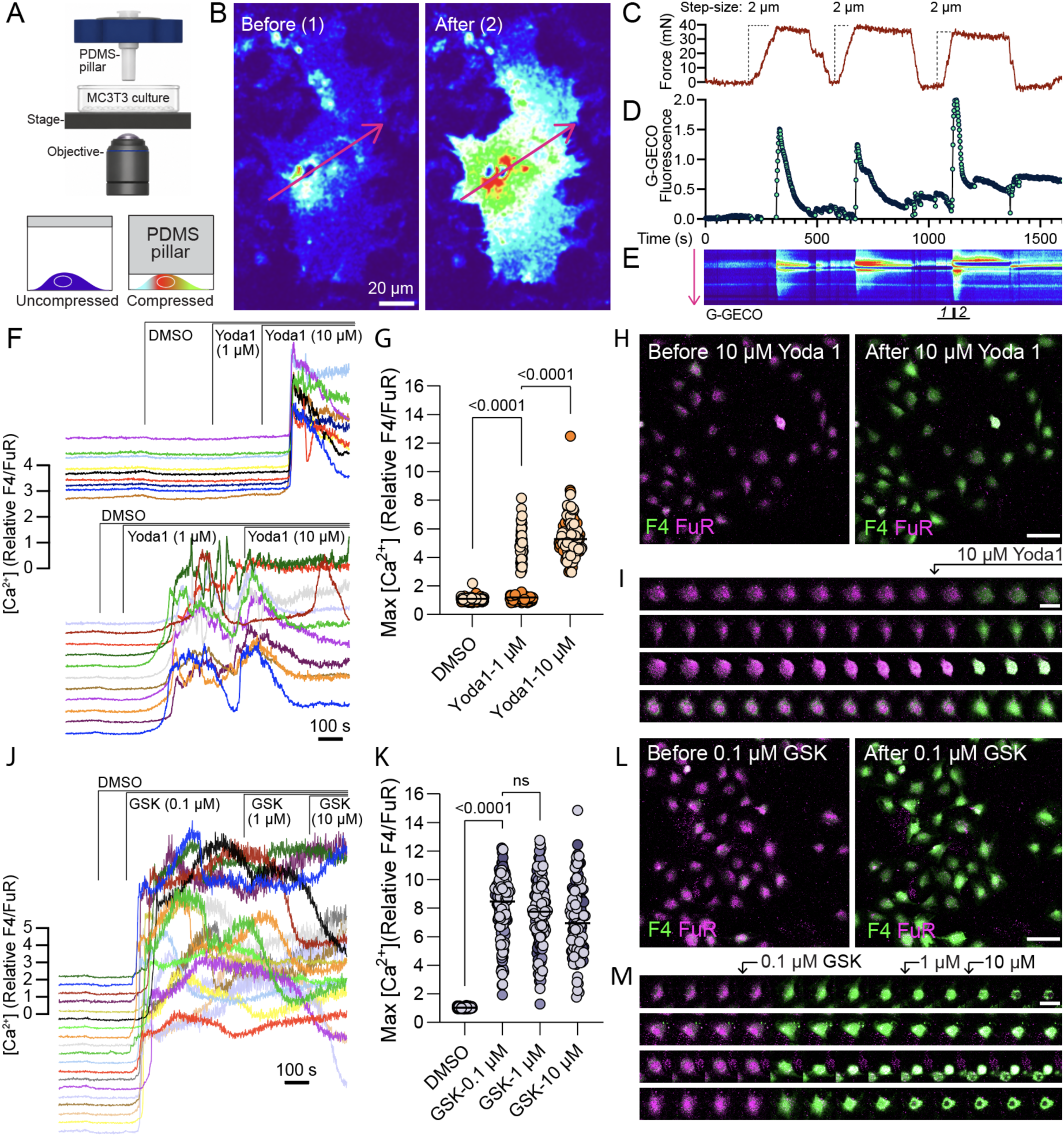
Sensitivity of MC3T3-E1 cells to mechanical compression and MIC agonists. **A.**Schematic of the Cell Press setup used to mechanically compress MC3T3-E1 cells in 2D culture during live imaging. The PDMS pillar is used to apply compression to cells loaded with a fluorescent Ca²⁺ reporter (G-GECO). **B.** Overview images of G-GECO fluorescence before (1) and after (2) compression in a single pre-osteoblastic MC3T3-E1 cell. The pink arrows in 1 and 2 indicate the linear profile plot used to construct the kymograph shown in E. **C.** Force measurements (mN) for a compression protocol in which an MC3T3-E1 was subjected to several compression and release cycles using incremental step sizes of 2 µm. The maximal force applied (∼40 mN) results from the cumulative compression effect of successive 2 ∝m displacement steps. **D.** Changes in cytosolic Ca^2+^ in an MC3T3-E1 over several compression cycle reported using the G-GECO reporter. **E.** Kymograph of the changes of the Ca^2+^ reporter fluorescence in response to compression over time across (pink arrow in B) an MC3T3-E1. The MC3T3-E1 in (B) is shown before (1) and after (2) the third compression. **F.** Changes in cytosolic Ca^2+^ (relative F4/FuR) in MC3T3-E1 cells before and after additions of DMSO, 1 μM Yoda1, 10 μM Yoda1. Representative traces from individual cells are shown from two independent experiment. **G.** Quantification of maximal calcium response (relative F4/FuR) in MC3T3-E1 cells after additions of DMSO, 1 μM Yoda1, 10 μM Yoda1. Black horizontal lines represent the median for each condition for three independent repeats (n=125 cells). Statistical analysis was performed using a Friedman test for paired data followed by a multiple comparison Dunn’s test. **H.** Fluorescence microscopy image (10X) of MC3T3-E1 stained with Fluo4 (F4, green) and FuraRed (FuR, magenta), before and after addition of 10 μM Yoda1 (scale bar 100 μm). **I.** Montage of fluorescence images of single cells stained with Fluo4 (green) and FuraRed (magenta), at regular interval throughout the time lapse imaging, before and after 10 μM Yoda1 addition (scale bar 50 μm). **J.** Changes in cytosolic calcium (relative F4/FuR) in MC3T3-E1 cells before and after additions of DMSO, 0.1 μM GSK1016790A, 1 μM GSK1016790A, 10 μM GSK1016790A. Representative traces from 10 out of 48 cells are shown for one experiment. **K.** Quantification of maximal calcium response (relative F4/FuR) in MC3T3-E1 cells after additions of DMSO, 0.1 μM GSK1016790A, 1 μM GSK1016790A, 10 μM GSK1016790A. Black horizontal lines represent the median for each condition for three repeats (n=135 cells). Statistical analysis was performed using a Friedman test for paired data followed by a multiple comparison Dunn’s test. **L.** Fluorescence microscopy image (10X) of MC3T3-E1 stained with Fluo4 (green) and FuraRed (magenta), before and after addition of 0.1 μM GSK1016790A (scale bar 100 μm). **M.** Montage of single cells stained with Fluo4 (green) and FuraRed (magenta), at regular interval throughout the time lapse imaging, before and after 0.1 μM GSK1016790A addition (scale bar 50 μm).

These observations demonstrate that our compression platform is compatible with monitoring MC3T3-E1 cell transduction of mechanical stimulation into intracellular calcium signaling. These responses are typically attributed to the activation of mechanosensitive ion channels (MICs), and so to confirm the activity of specific MICs in the MC3T3-E1 cells we exposed them to small molecule agonists of Piezo1 (Yoda1) or TRPV4 (GSK1016790A). Ca^2+^ responses were monitored using the ratiometric fluorescence readout Fluo-4/Fura Red (F4/FuR), where increases in the ratio reflect elevated intracellular Ca²⁺ levels. Dimethyl sulfoxide (DMSO) was added as a negative control stimulation, both as it is solvent used in the Yoda1 and GSK1016790A solutions, and to ensure that the flow rate for reagents added to the dish was not sufficient to induce a shear-based mechanical response. Stimulation with 1 µM Yoda1 resulted in variable responses, with Ca²⁺ elevation detected in only one of the three replicates (Fig. 2F, examples in lower traces). In contrast, 10 µM Yoda1 consistently triggered Ca²⁺ responses (Fig. 2F-H). MC3T3-E1 cells exhibited sensitivity to 0.1 ∝M of the TRPV4 channel agonist GSK1016790A, evident as a rapid increase in cytosolic Ca^2+^. Subsequent additions of 1 ∝M and 10 ∝M GSK1016790A resulted in sustained Ca^2+^ elevations or gradual decreases. However, on average the Ca^2+^ response to 0.1 ∝M represented the maximum signal induced during all treatments. (Fig. 2J and K). GSK1016790A-induced increases in Ca^2+^ were also occasionally accompanied by distinct morphological changes, including cell contraction, and Ca^2+^ influx via TRPV4 has previously implicated in actomyosin-mediated contractility^38^ (Figure 2 L, M). MC3T3-E1 pre-osteoblastic cells are commonly used as an osteoblast model and have previously been reported to express both Piezo1^39–41^ and TRPV4^40,42,43^. Piezo1 has also previously been implicated in MC3T3-E1 cells electrophysiological mechanoresponses to poking^39^. Other studies have additionally suggested an important role for TRPV4 in calcium responses induced by shear stress^40^, which is experienced by osteocytes arising from load-induced flow in the fluid filled lacunar-canalicular network of bone tissues^44^.

Here, we demonstrated the applicability of the Cell Press platform for studying mechanoresponses in MC3T3-E1 cells under controlled compressive loading. We confirm the presence of functional Piezo1 and TRPV4 MICs in the MC3T3-E1 cells, indicating that they are likely among the mechanosensors contributing to the Ca^2+^ changes induced by the Cell Press in Fig. 2.

### Melt-electrowritten PCL scaffold characterization and assembly with the Cell Press for compression

Porous PCL scaffolds comprised of 90 layers of overlapping grids were fabricated using melt electrowriting (MEW). The resulting scaffold architecture exhibited the intended square pore geometry, observed using a stereo– or confocal microscope equipped with differential interference contrast (Fig. 3A, B). The printing parameters produced uniform microfibers with an average diameter of 19.6 ± 2.9 μm (Fig. 3C). The intended interfiber space was 300 μm, and the measured spacing was 274.5 ± 8.7 μm (Fig. 3D). This reduction is likely attributable to interfiber attraction effects, known to occur during the MEW process^4^.

**Figure 3.**
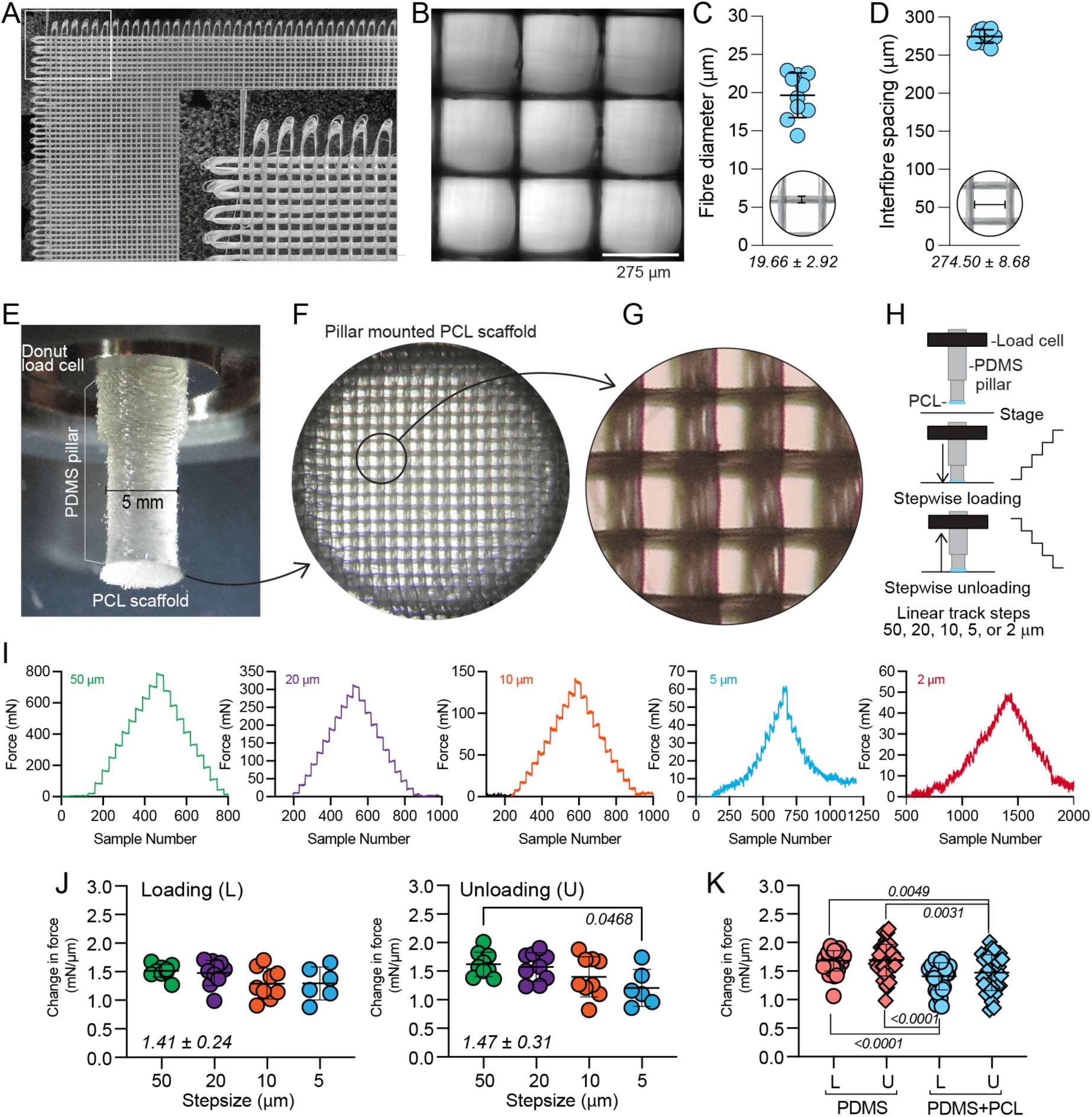
Characterization of the melt-electrowritten PCL scaffolds and assembly with the Cell Press. **A**. Stereomicroscopy images of PCL scaffold (90 layers) designed with an intended interfiber spacing of 300 µm. Insert illustrates the looping at the end of each linear horizontal and vertical fiber path. B. Transmitted light images of 3 x 3 pores of a PCL scaffold imaged by confocal microscopy with a Plan-Apochromat 10x/0.45 objective. C. Quantitative assessment of the average interfiber space (μm) in the PCL scaffolds from n = 10 measurements (mean ± SD values stated). D. Quantitative assessment of the average PCL fiber diameter (μm) from n = 10 measurements (mean ± SD values stated). E. Photograph of a 90-layer PCL scaffold fixed to the surface of the PDMS pillar, which is mounted in the donut load cell of the cell compression tool. F. Stereomicroscope image of a 90-layer PCL scaffold on the PDMS pillar. G. High magnification view of the scaffold in F illustrating the multiple layers of PCL fibers. H. Overview of the protocol applied to load and unload the PDMS pillar with the PCL scaffold attached (PDMS + PCL), directly against the microscope stage. I. Dynamic force measurements were acquired at a rate of 1 sample/second with the Futek donut load cell interfaced with a USB signal conditioner, with signals visualized and logged using Sensit data acquisition software. Each panel represent the recorded forces (mN) during PDMS pillar loading and unloading for each of the indicated step sizes (50 μm to 2 μm). **J.** Quantification of the average change in force (mN)/ μm, calculated for each of the indicated step sizes (50 μm – 5 μm) during the loading phase (left plots) and unloading phase (right plots). The average change in force (mN/μm) + standard deviation for all quantified step sizes is presented in the respective plots. **K.** Comparison of average changes in force/displaced mm during loading (L) and unloading (U) for all step sizes with the PDMS alone (PDMS) (from Fig. 1), and for the PCL scaffold mounted on the PDMS pillar. Statistically significant differences were assessed using a one-way ANOVA with multiple comparisons. P-values are stated in the figure panels.

To investigate compressive forces applied through the PCL scaffold, it was mounted onto the PDMS pillar (Fig. 3E-G) as follows. A circular disc was excised from the scaffold using a biopsy punch, and was affixed to the exposed surface of the pillar. Inspection of the scaffold confirmed that its architecture remained intact following the mounting process (Fig. 3F-G). The compression performance of the PCL-mounted PDMS pillar was evaluated through force measurements during loading and unloading cycles as described in Fig. 1 (Fig. 3H). The compressive force response of the PDMS–PCL system was recorded across step sizes down to 2 μm (Fig. 3I). Actuating the linear track with step sizes ranging from 50 ∝m to 5 μm generated a force (mN) per displacement micrometer of 1.41 ± 0.24 mN/μm during loading and 1.47 ± 0.31 mN/μm during unloading. This was consistent for all step-sizes during loading, but a slight decrease was observed between 50 ∝m and 5 ∝m steps during the unloading phase (Fig. 3J). This establishes that the PCL scaffold can be used to exert reproducible force across incremental displacements, with comparable force– displacement behavior observed across the tested step sizes in loading and unloading cycles. In comparison to the PDMS pillar alone, the PDMS–PCL system exhibited a moderate reduction in the average change in force per displaced micrometer (Fig. 3K). This is likely attributed to the PCL scaffold structure accommodating a component of the compressive force. Notably, the measured forces reflect the combined mechanical behavior of the PDMS pillar and the scaffold, and do not directly represent the intrinsic mechanical properties of the scaffold alone. Therefore, it is not intended to replace conventional mechanical testing systems for the characterization of biomaterials or scaffolds properties under compression, as typically performed using universal testing machines^45,46^. Collectively, these results establish successful integration of the biomaterial scaffold into the Cell Press platform and demonstrate dynamic, low-micrometre-resolution control of uniaxial displacement alongside real-time quantification of compressive forces.

### Imaging MC3T3-E1 Ca^2+^ mechanoresponses to PCL scaffold-transmitted compression

The capacity of the PCL scaffold to transmit compression-induced mechanoresponses was evaluated by live imaging cytosolic Ca^2+^ dynamics in a monolayer of MC3T3-E1 cells during a series of compression events. Contact with the cells was confirmed by an increase in the measured force profiles, and were aligned with single-cell Ca^2+^ measurements to detect overlap between compression and Ca^2+^ signaling events. For quantification, maximum Ca^2+^ responses from individual cells when uncompressed (baseline) were compared with maximum responses during compression phases. Initial experiments were conducted with the PDMS pillar alone (Fig. 4A–C). Representative traces reveal transient responses during sustained compression and heterogeneity in cellular responses across the population, with not all cells responding to all compression events (Fig. 4A, and kymograph in 4C). Nonetheless, a significant increase in the average maximum cytosolic Ca^2+^ levels was observed during compression (Fig. 4B). The observed variability in responses likely reflects differences in z-axis height of the cells, as ‘taller’ cells will come in contact with the descending pillar surface before ‘shorter’ neighbors. Additionally, the degree of cell adhesion with the underlying substrate may alter plasma membrane tension, which in turn could modulate stretch-activated MIC responses^47,48^. Further, intercellular differences in the relative expression levels of MICs may also contribute to defining a cells potential to respond to a given mechanical stimulus. Finally, it is relevant to consider the horizontal alignment of the pillar surface relative to the MC3T3-E1 monolayer as an additional contributing factor to this variability. The 3D-printed part that holds the load cell and PDMS pillar is inserted into the arm of the device that is in turn attached to the piezoelectric track (Fig. 1B). This part can be freely rotated within the arm, which is intended to facilitate aligning the surface of the pillar parallel with the surface of the culture dish. However, while the precision of this alignment is aided by viewing the pillar surface through the microscope, the adjustment is manual and variability in observed responses may result from the pillar’s surface not being aligned in parallel to the cell layer.

**Figure 4.**
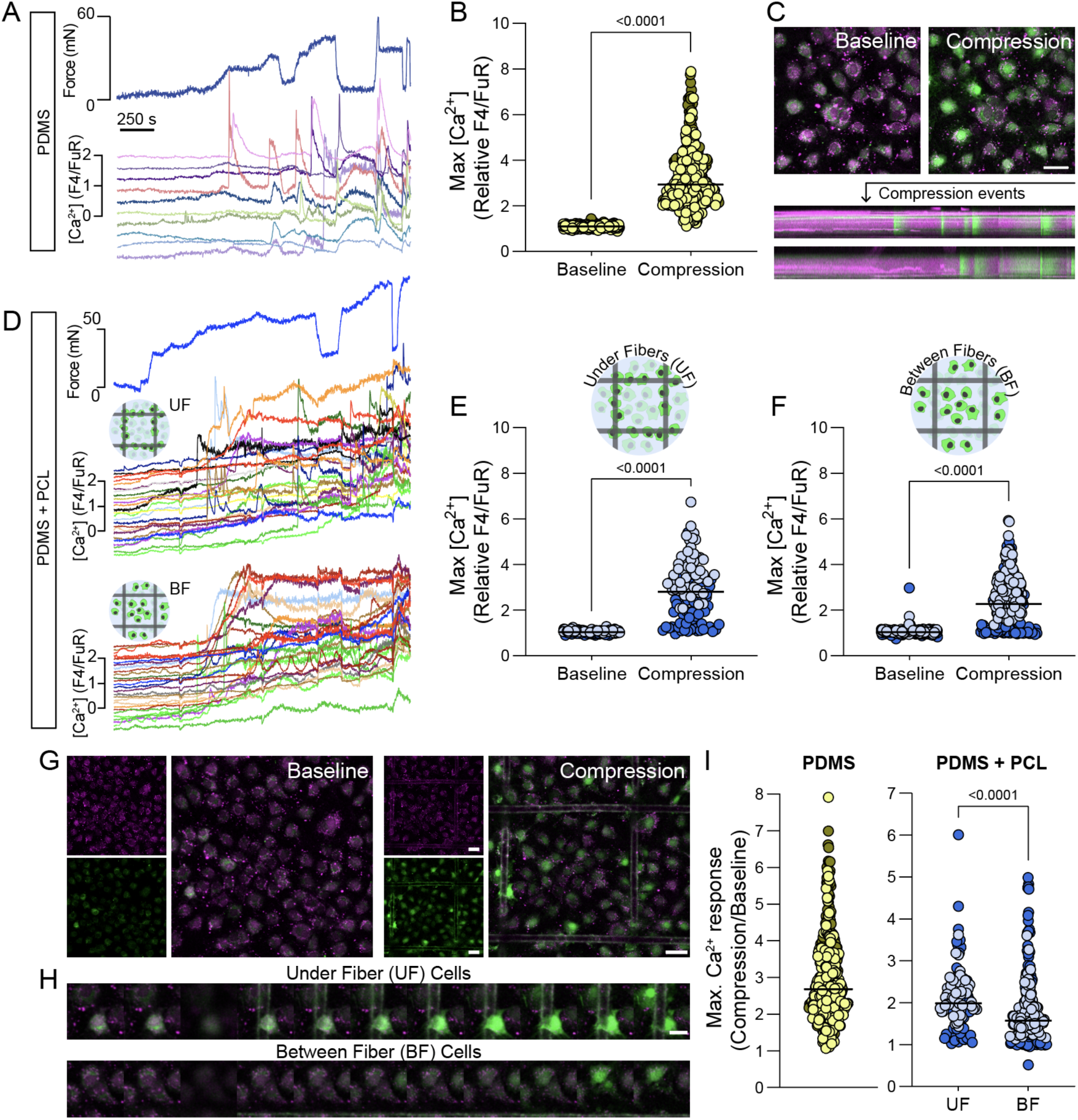
Platform to investigate cellular responses to mechanical stimulation using PCL scaffolds. **A**. Force measurements (mN) recorded during a compression protocol in which pre-osteoblastic MC3T3-E1 cells were subjected to repeated compression–release cycles using a PDMS pillar, together with the corresponding changes in cytosolic calcium (relative F4/FuR). Arrows indicate the onset of each compression (C). Representative traces from 10 out of 186 cells are shown from one experiment. **B.** Quantification of maximal calcium responses (relative F4/FuR) in MC3T3-E1 cells at baseline (before compression) and after compression. The maximal calcium response across all compression cycles was extracted for each cell. Black horizontal lines indicate medians for each condition across two independent experiments (n = 374 cells). Statistical analysis was performed using an unpaired, non-parametric Wilcoxon test. **C.** Fluorescence microscopy images (10×) of MC3T3-E1 cells stained with Fluo-4 (green) and Fura Red (magenta) at baseline and after compression (scale bar: 200 µm). Kymographs show temporal changes in Ca²⁺ reporter fluorescence during compression in two representative cells. **D.** Cytosolic Ca²⁺ responses (relative F4/FuR) during repeated compressions using a PCL scaffold attached to the PDMS pillar. Responses are shown for cells located under the scaffold fibers and for cells situated between the fibers. Arrows mark compression onset (C). Representative traces from 10 out of 118 cells (under fibers) and 10 out of 54 cells (between fibers) are shown. **E.** Quantification of maximal calcium responses (relative F4/FuR) for MC3T3-E1 cells located under PCL scaffold fibers at baseline and after compression. The maximal calcium response across all compression cycles was extracted for each cell. Black horizontal lines indicate medians across two independent experiments (n = 248 cells). Statistical analysis was performed using a paired t-test. **F.** Quantification of maximal calcium responses (relative F4/FuR) for MC3T3-E1 cells located between PCL scaffold fibers at baseline and after compression. The maximal calcium response across all compression cycles was extracted for each cell. Black horizontal lines indicate medians across two independent experiments (n = 108 cells). Statistical analysis was performed using a matched, non-parametric Wilcoxon test. **G.** Fluorescence microscopy images (10×) of MC3T3-E1 cells stained with Fluo-4 and Fura Red at baseline and after compression with the PCL scaffold (scale bar: 200 µm). **H.** Montage showing two representative cells located under PCL scaffold fibers and two representative cells positioned between fibers during compression. I. Fold change in maximal calcium response relative to baseline for cells compressed with the PDMS pillar alone and with the PDMS pillar in combination with a PCL scaffold. Comparisons are shown for cells under fibers (UF) and between fibers (BF). Statistical analysis was performed using a non-parametric, unpaired Mann–Whitney test.

Next, compression-induced signaling events were investigated with the Cell Press-mounted PCL scaffolds. Cells were spatially categorized based on whether they were positioned directly beneath and physically in contact with a PCL fiber during compression, or located in the interfiber porous spaces. Increases in cytosolic Ca^2+^ were observed for both cell categories across multiple compression events (Fig. 4D), and the maximum Ca^2+^ responses recorded in individual cells were significantly increased relative to those measured in uncompressed baseline recordings (Fig. 4E-H). Nonetheless, significantly higher maximum Ca^2+^ responses were recorded for cells directly under the scaffold fibers than for cells in the interfiber spaces (Fig. 4I). Interestingly, the induction of Ca^2+^ responses in cells located in between fibers suggests that mechanisms other than direct mechanical compression contribute to the signal propagation observed throughout the MC3T3-E1 monolayer.

Several factors may contribute to these apparently indirect mechanoresponses. In confluent MC3T3-E1 cultures, the establishment of cell–cell contacts including gap junctions may facilitate propagation of Ca^2+^ signals from a compressed cell to a non-compressed neighbor^49^. Additionally, such cell-cell contacts would create a physical connection between plasma membranes that could mediate the transmission of membrane tension changes between cells. It is also possible that mechanically induced Ca^2+^ signaling triggers paracrine pathways, leading to Ca^2+^ increases in neighboring cells via for example ATP release. Extracellular ATP can activate purinergic receptors, resulting in Ca²⁺ mobilization from intracellular stores such as the endoplasmic reticulum, which has previously been implicated as the predominant mechanism of intercellular Ca^2+^ signaling in MC3T3-E1 cells^50^. Finally, compression with the PCL scaffold may induce changes in local fluid flow patterns in the scaffold pores, which could induce shear stress-triggered Ca^2+^ mechanoresponses^51^.

In other studies, compression has been applied to cell-seeded scaffolds to provide external mechanical cues, mimic aspects of load-bearing conditions, and evaluate cellular responses through downstream analyses of gene and protein expression^20,52,53^. The Cell Press platform enables real-time observation of dynamic and transient cellular responses to mechanical stimulation. The method could be further extended to more complex configurations, by studying Ca^2+^ responses in cells seeded directly on the scaffolds, and methods for attaching cells to these PCL scaffolds have been established^26^. The platform can also be applied for mechanistic studies of the role of mechanosensitive ion channels, such as Piezo1 and TRPV4, in scaffold-transmitted compression signaling.

### Conclusion

This study establishes the feasibility of integrating biomaterial scaffolds with the Cell Press platform for monitoring dynamic and transient mechanosensitive Ca^2+^ responses to compressive loading and unloading. A specific insight revealed by this approach is that the porous PCL scaffolds elicited direct and indirect MC3T3-E1 Ca^2+^ responses to scaffold-transmitted compression. Dynamic interactions between engineered scaffolds and relevant mechanosensing cell populations remain relatively underexplored, but may help inform scaffold designs to potentially improve *in vivo* function. The described Cell Press-based approach represents a promising strategy to investigate cellular responses to scaffold-transmitted mechanical stimulation, with potential relevance for implants in load-bearing tissues, including cartilage and bone, where physiological forces may be transmitted across the scaffold-tissue interface.

## Materials and Methods

### Cell Press tool assembly and operation

The Cell Press is a custom-built compression device composed of a linear piezoelectric stage (SmarAct GmbH, Oldenburg, Germany), a Futek LTH300 donut load cell (Microepsilon, Sensotest AB, Järfälla, Sweden), and a flexible polydimethylsiloxane (PDMS) pillar. The PDMS pillar was fabricated using Sylgard 184 silicone elastomer kit (Sigma-Aldrich Sweden AB, Stockholm, Sweden) at a 10:1 base-to-curing-agent ratio and cast using a custom mold produced with a Form3 3D printer (Formlabs)^24^. The PDMS pillar was designed to be fitted in the load cell, the inner surface of which is load sensing. The components were assembled using 3D-printed parts that enable mounting of the piezoelectric track onto the microscope stage and connect the track to the load cell containing the PDMS pillar. The piezoelectric stage is controlled via an MCS2 manual controller (SmarAct), allowing stepwise displacement of the PDMS pillar. The load cell was connected via a USB signal conditioner (Futek) to a PC running Sensit software for real-time force acquisition (Futek).

### Melt-electrowriting of poly(ε-caprolactone) scaffolds

Poly(ε-caprolactone) (PCL) was melt-electrowritten using a custom-built melt electrowriting device and printing protocol optimized^26^ at ETH Zürich in the Institute for Biomechanics. Medical-grade PCL (PURASORB PC 12, Corbion Inc., Netherlands) was extruded from a syringe maintained above its melting temperature (80 °C) within the melt head, through a 22G needle (Nordson, Westlake, OH, USA) at a pressure of 1 bar onto a translating collector. The porous scaffold architecture was defined using a custom Python script generating G-code to control collector motion. Fiber wall spacing was set to 300 μm, with successive layers deposited at 90° relative to each other to produce a square pore geometry. The process began at a nozzle-to-collector distance of 3.6 mm and an applied voltage of 5.4 kV, with the distance incremented by 20 µm per layer. To compensate for insulation effects from previously deposited layers, the applied voltage was increased by 10 V after each layer. The scaffold dimensions were specified as 20 x 20 mm (width × length) with a total of 90 layers.

### PCL scaffold fiber characterization and attachment to the Cell Press

The PCL scaffolds were imaged using a stereomicroscope and a confocal microscope LSM 700 (Zeiss, Germany). Filament diameter and fiber spacing were measured on Image J (Fiji) in ten different locations of a scaffold using a Differential Interference Contrast (DIC) image captured on the confocal microscope with a Plan-Apochromat 10×/0.45 (Zeiss) objective. Six mm diameter discs were excised from PCL scaffolds using a biopsy punch and were attached to the surface of the PDMS compression pillar using a water soluble, polyvinylacetate-based liquid glue..

### Stepwise loading and unloading

Force measurements (mN) during loading and unloading using different linear track step sizes, were performed on the solid microscope stage. The PDMS pillar, or the PCL scaffold, were brought into near contact with the microscope stage surface prior to force recording. Force measurements were tared to zero before each loading experiment. Displacement steps (µm) were defined using the MCS2 manual controller and applied at regular 30-second intervals. Unloading was performed by reversing the displacement until the pillar or scaffold was no longer in contact with the stage.

### Cell culture

The MC3T3-E1 subclone 4 pre-osteoblastic cell line (CRL-2593, ATCC) was used for compression testing. Cells were cultured using alpha-MEM supplemented with 10% FBS and 1% Penicillin/Streptomycin (Thermo Fisher Scientific, Sweden), referred to as cell culture medium, under standard incubation conditions of 37 °C, 5% CO_2_.

### Staining reagents and MIC agonists

The calcium indicators Fluo-4 and Fura Red (Thermo Fisher Scientific, Uppsala, Sweden) were suspended to 1 mM with dimethylsulfoxide (DMSO; ThermoFisher Scientific, Uppsala, Sweden). Stock solutions of the Piezo1 agonist Yoda1 (Tocris, Bio-Techne, Bristol, UK) were diluted in DMSO to 10 mM, and the TRPV4 agonist GSK1016790A (Tocris) was diluted in DMSO to 50 mM.

### Live cell calcium imaging using Fluo-4 and Fura Red

MC3T3-E1 were seeded in glass-bottom culture dishes (µ-Dish 35 mm, high, Ibidi GmbH) and cultured in cell culture media. Cells were incubated with Ca^2+^ reporting dyes Fluo-4 and Fura red diluted to 5 μM in phenol red free Opti-MEM (Thermo Fisher Scientific, Sweden) for at least 1 h prior to imaging. Cells were washed with Opti-MEM before imaging. Time-lapse imaging of Fluo-4 (F4) and Fura Red (FuR) fluorescence channels was performed on an LSM700 confocal microscope (Zeiss, Germany) equipped with an incubation chamber (37 °C), using a Plan-Apochromat 20x/0.8 (Zeiss) objective. F4 and FuR were excited at 488 nm. Signals were acquired simultaneously using two independent photomultiplier tube (PMT) detectors. Before stimulation (mechanical compression or MIC agonists), a baseline of Fluo-4 and Fura Red fluorescence was recorded for several minutes.

#### Mechanical stimulation

For compression experiments, cells were seeded at a density of 28,570 cells/cm² (100,000 cells/dish) and imaged 2–3 days after seeding. Prior to stimulation, a reference dish without cells or medium was used to align the PDMS pillar with the dish surface and to set the zero position for the z-axis height of the pillar, to determine the position at which the pillar contacts the cell layer. Force measurements were calibrated by taring to 0 mN before compression. Force recording (via the Sensit software) was initiated simultaneously with fluorescence imaging, and acquired at a rate of 1 measurement/sec. A stable baseline force signal was recorded for several minutes before applying compression. Mechanical compression was applied using step sizes of 2 µm, 1 µm, and 200 nm. The focal plane was adjusted when needed during compression experiments to compensate for any displacement of the culture dish. The timing of compression events was recorded by noting the start of compression and end of compression frames of the time-lapse recordings, and this was later aligned with the Ca^2+^ recordings.

#### Yoda1 and GSK1016790A stimulation

For experiments with MIC agonists, cells were seeded at 5,700 cells/cm² (20,000 cells/dish) and imaged 2 days post-seeding. Yoda1 stock solutions were diluted to 10 μM and 1 μM in Opti-MEM. GSK1016790A stock solutions were diluted to 10 μM, 1 μM and 0.1 μM in Opti-MEM. During imaging, DMSO (control), Yoda1, or GSK1016790A solutions were added directly to the culture dish via a syringe-based delivery system. A 1 mL syringe (BD Plastipak) fitted with a 0.55 mm × 25 mm needle (BD Microlance 3) was connected to 45 cm of tubing (inner diameter 0.51 mm; Masterflex® Transfer Tubing, Tygon ND-100-80, VWR). The tubing was inserted into the culture dish through a custom-designed 3D-printed cap, enabling reagent addition without interrupting imaging. All solutions were prepared to account for dilution effects resulting from cumulative additions to the imaging volume. The timing for the addition of control or MIC agonist solutions was recorded by noting the frame numbers of the time-lapse recordings at the start and end of the injection.

### Live calcium imaging using G-GECO

Adenoviral particles (E5 serotype, > 10^12^–10^13^ virus particles/ml) carrying 5HT6-G-GECO. This construct is under CMV-promoter control and encodes serotonin receptor 6, which facilitates anchorage of the fused calcium-sensitive fluorescent reporter (GECO) to the plasma membrane (Vector Biolabs). Cells were incubated with 2 ∝l of stock virus for 3 hours at 37 °C and then transferred back to fresh culture medium, expression was maintained for at least 2 days before performing imaging experiment^25^.

### Image analysis and processing

Time-lapse sequences of Fluo-4 and Fura Red fluorescence were analyzed using the Fiji format of ImageJ. Single cells in the field of view were segmented using the Cellpose Python algorithm (https://www.cellpose.org/). Each cell was added as a region of interest (ROI) to the ROI manager using Fiji software^54^. Fluo-4 and Fura Red fluorescence intensity were measured for each cell for the duration of the timelapse. The Fluo-4/Fura red ratio was expressed to investigate calcium flux during compression events or MIC agonist exposure. Fluor-4/Fura red ratio for each timepoint of the time lapse was normalized to the ratio at the first timepoint, permitting a relative comparison of changes in calcium dynamics between all cells throughout the timeseries. For compression with the PCL scaffolds, segmented cells were categorized according to whether they were observed to be under the fibers or between the fibers of the PCL scaffold during the compression events.

### Compression events analysis

The real-time force recordings collected using the Sensit software were manually aligned with the corresponding Fluo-4/Fura Red Ca^2+^ responses for the experiment’s duration. The maximum Fluo-4/Fura Red responses detected across all compression events for a given experiment were identified for each cell and presented as scatter plots compared to the baseline maximal response (before compression). Statistical analyses were performed using GraphPad Prism (version 10.16.1), and the specific statistical tests applied are indicated in the figure legends.

## Acknowledgments

The authors acknowledge funding provided by the European Union’s Horizon 2020 Research and Innovation Programme to the project BioTriB, grant agreement No. 956004. This study was conducted as part of the Additive Manufacturing for the Life Sciences (AM4Life) consortium, which is funded by Sweden’s Innovation Agency Vinnova (grant number 2019-00029), and by additional grants from the Swedish Cancer Society (Cancerfonden; grant numbers 20 1285 PjF and 23 2692 Pj 01 H), the Göran Gustafsson’s Foundation (Göran Gustafssons Stiftelser), and a G. Bergmark travel scholarship. 3D printing was performed at U-PRINT: Uppsala University’s 3D-printing facility at the Disciplinary Domain of Medicine and Pharmacy and SciLifeLab Uppsala. We would like to thank and acknowledge Dr. Gonzalo Manuel Sanchez, Dept. of Medical Cell Biology, Uppsala University for providing G-GECO expressing MC3T3-E1 cell cultures.

## Declaration of AI use

The authors acknowledge the use of generative artificial intelligence (AI) tools, including OpenAI ChatGPT (GPT-5.5 release) and Microsoft Copilot for language editing and manuscript refinement. All scientific content, interpretations, analyses, and conclusions in the final manuscript are the authors own.

## Data availability

All data supporting the findings of this study are available on reasonable request to the corresponding author.

## Notes

### Competing Interest Statement

The authors have declared no competing interest.

## References

1. Sanchez-Adams, J., Leddy, H. A., McNulty, A. L., O’Conor, C. J. & Guilak, F. The Mechanobiology of Articular Cartilage: Bearing the Burden of Osteoarthritis. Curr Rheumatol Rep 16, 451 (2014).

2. Vinatier, C. & Guicheux, J. Cartilage tissue engineering: From biomaterials and stem cells to osteoarthritis treatments. Annals of Physical and Rehabilitation Medicine 59, 139–144 (2016).

3. Loewner, S. et al. Recent advances in melt electro writing for tissue engineering for 3D printing of microporous scaffolds for tissue engineering. Front. Bioeng. Biotechnol. 10, 896719 (2022).

4. O’Neill, K. L. & Dalton, P. D. A Decade of Melt Electrowriting. Small Methods 7, 2201589 (2023).

5. Zhou, S. et al. Melt Electrowriting: A Promising 3D Printing Technology for Cartilage and Osteochondral Repair. Advanced Therapeutics 7, 2300218 (2024).

6. Jangi, R., Zgeib, R., Zaeri, A. & Chang, R. C. Melt electrowriting for bone repair: A review of scaffold architectures and material selection. Bioprinting 55, e00474 (2026).

7. Vogel, V. & Sheetz, M. Local force and geometry sensing regulate cell functions. Nat Rev Mol Cell Biol 7, 265–275 (2006).

8. Zarka, M., Haÿ, E. & Cohen-Solal, M. YAP/TAZ in Bone and Cartilage Biology. Front Cell Dev Biol 9, 788773 (2022).

9. Di, X. et al. Cellular mechanotransduction in health and diseases: from molecular mechanism to therapeutic targets. Sig Transduct Target Ther 8, 282 (2023).

10. Heller, S. & O’Neil, R. G. Molecular Mechanisms of TRPV4 Gating. in TRP Ion Channel Function in Sensory Transduction and Cellular Signaling Cascades (CRC Press/Taylor & Francis, 2007).

11. Coste, B. et al. Piezo1 and Piezo2 Are Essential Components of Distinct Mechanically Activated Cation Channels. Science 330, 55–60 (2010).

12. Liu, Y. et al. Mechanosensitive channel Piezo1 in calcium dynamics: structure, function, and emerging therapeutic strategies. Front. Mol. Biosci. 12, 1693456 (2025).

13. Berridge, M. J., Bootman, M. D. & Roderick, H. L. Calcium signalling: dynamics, homeostasis and remodelling. Nat Rev Mol Cell Biol 4, 517–529 (2003).

14. Clapham, D. E. Calcium Signaling. Cell 131, 1047–1058 (2007).

15. Zayzafoon, M. Calcium/calmodulin signaling controls osteoblast growth and differentiation. J. Cell. Biochem. 97, 56–70 (2006).

16. McNulty, A. L., Leddy, H. A., Liedtke, W. & Guilak, F. TRPV4 as a Therapeutic Target for Joint Diseases. Naunyn Schmiedebergs Arch Pharmacol 388, 437–450 (2015).

17. Zhou, T., et al. Piezo1/2 mediate mechanotransduction essential for bone formation through concerted activation of NFAT-YAP1-ß-catenin. eLife 9, e52779 (2020).

18. Wang, L., et al. Mechanical sensing protein PIEZO1 regulates bone homeostasis via osteoblast-osteoclast crosstalk. Nat Commun 11, 282 (2020).

19. Ehrlich, P. J. & Lanyon, L. E. Mechanical Strain and Bone Cell Function: A Review. Osteoporos Int 13, 688–700 (2002).

20. Rath, B., Nam, J., Knobloch, T. J., Lannutti, J. J. & Agarwal, S. Compressive forces induce osteogenic gene expression in calvarial osteoblasts. Journal of Biomechanics 41, 1095–1103 (2008).

21. Charras, G. T., Lehenkari, P. P. & Horton, M. A. Atomic force microscopy can be used to mechanically stimulate osteoblasts and evaluate cellular strain distributions. Ultramicroscopy 86, 85–95 (2001).

22. Lomakin, A. J., et al. The nucleus acts as a ruler tailoring cell responses to spatial constraints. Science 370, eaba2894 (2020).

23. Onal, S., Alkaisi, M. M. & Nock, V. Microdevice-based mechanical compression on living cells. iScience 25, 105518 (2022).

24. O’Callaghan, P., et al. Piezo1 activation attenuates thrombin-induced blebbing in breast cancer cells. Journal of Cell Science 135, jcs258809 (2022).

25. Sanchez, G. M., et al. The β-cell primary cilium is an autonomous Ca2+ compartment for paracrine GABA signaling. J Cell Biol 222, e202108101 (2022).

26. Santschi, M. X. T., et al. Mechanical and Biological Evaluation of Melt-Electrowritten Polycaprolactone Scaffolds for Acetabular Labrum Restoration. Cells 11, 3450 (2022).

27. Miranda, I., et al. Properties and Applications of PDMS for Biomedical Engineering: A Review. J Funct Biomater 13, 2 (2021).

28. Srivastava, N., Kay, R. R. & Kabla, A. J. Method to study cell migration under uniaxial compression. MBoC 28, 809–816 (2017).

29. Andolfi, L., et al. Planar AFM macro-probes to study the biomechanical properties of large cells and 3D cell spheroids. Acta Biomaterialia 94, 505–513 (2019).

30. Aung, A., Davey, S. K., Theprungsirikul, J., Kumar, V. & Varghese, S. Deciphering the mechanics of cancer spheroid growth in 3D environments through microfluidics driven mechanical actuation. Adv Healthc Mater 12, e2201842 (2023).

31. Pandey, M., et al. Viscoelastic properties of tumor spheroids revealed by a microfluidic compression device and a modified power law model. Soft Matter 22, 1618–1629 (2026).

32. Kim, T. K. & Jeong, O. C. Intracellular calcium-expression-display (ICED) device operated by compressive stimulation of cells. Microelectronic Engineering 98, 703–706 (2012).

33. Lee, D., Erickson, A., You, T., Dudley, A. T. & Ryu, S. Pneumatic microfluidic cell compression device for high-throughput study of chondrocyte mechanobiology. Lab Chip 18, 2077–2086 (2018).

34. Onal, S., Alkaisi, M. M. & Nock, V. A Flexible Microdevice for Mechanical Cell Stimulation and Compression in Microfluidic Settings. Front. Phys. 9, 654918 (2021).

35. Schriefer, J. L., Warden, S. J., Saxon, L. K., Robling, A. G. & Turner, C. H. Cellular accommodation and the response of bone to mechanical loading. Journal of Biomechanics 38, 1838–1845 (2005).

36. Turner, C. H. Toward a Mathematical Description of Bone Biology: The Principle of Cellular Accommodation. Calcif Tissue Int 65, 466–471 (1999).

37. Lewis, A. H., Cui, A. F., McDonald, M. F. & Grandl, J. Transduction of Repetitive Mechanical Stimuli by Piezo1 and Piezo2 Ion Channels. Cell Reports 19, 2572–2585 (2017).

38. Gilchrist, C. L., et al. TRPV4-mediated calcium signaling in mesenchymal stem cells regulates aligned collagen matrix formation and vinculin tension. Proceedings of the National Academy of Sciences 116, 1992–1997 (2019).

39. Sun, W., et al. The mechanosensitive Piezo1 channel is required for bone formation. eLife 8, e47454 (2019).

40. Yoneda, M., et al. PIEZO1 and TRPV4, which Are Distinct Mechano-Sensors in the Osteoblastic MC3T3-E1 Cells, Modify Cell-Proliferation. International Journal of Molecular Sciences 20, 4960 (2019).

41. Zhang, G., Li, X., Wu, L. & Qin, Y.-X. Piezo1 channel activation in response to mechanobiological acoustic radiation force in osteoblastic cells. Bone Res 9, 16 (2021).

42. Mizoguchi, F., et al. Transient receptor potential vanilloid 4 deficiency suppresses unloading-induced bone loss. Journal of Cellular Physiology 216, 47–53 (2008).

43. Li, Y., Yang, Y., Wang, X., Li, L. & Zhou, M. Extracellular osmolarity regulates osteoblast migration through the TRPV4-Rho/ROCK signaling. Commun Biol 8, 515 (2025).

44. Murshid, S. A. Bone permeability and mechanotransduction: Some current insights into the function of the lacunar-canalicular network. Tissue and Cell 75, 101730 (2022).

45. Abbasi, N., Abdal-hay, A., Hamlet, S., Graham, E. & Ivanovski, S. Effects of Gradient and Offset Architectures on the Mechanical and Biological Properties of 3-D Melt Electrowritten (MEW) Scaffolds. ACS Biomater. Sci. Eng. 5, 3448–3461 (2019).

46. Liu, H., Ahlinder, A., Yassin, M. A., Finne-Wistrand, A. & Gasser, T. C. Computational and experimental characterization of 3D-printed PCL structures toward the design of soft biological tissue scaffolds. Materials & Design 188, 108488 (2020).

47. Yao, M., et al. Force– and cell state–dependent recruitment of Piezo1 drives focal adhesion dynamics and calcium entry. Sci. Adv. 8, eabo1461 (2022).

48. Cheng, D., Wang, J., Yao, M. & Cox, C. D. Joining forces: crosstalk between mechanosensitive PIEZO1 ion channels and integrin-mediated focal adhesions. Biochemical Society Transactions 51, 1897–1906 (2023).

49. Xia, S.-L. & Ferrier, J. Propagation of a calcium pulse between osteoblastic cells. Biochemical and Biophysical Research Communications 186, 1212–1219 (1992).

50. Huo, B., Lu, X. L., Costa, K. D., Xu, Q. & Guo, X. E. An ATP-dependent mechanism mediates intercellular calcium signaling in bone cell network under single cell nanoindentation. Cell Calcium 47, 234–241 (2010).

51. Kapur, S., Baylink, D. J. & William Lau, K.-H. Fluid flow shear stress stimulates human osteoblast proliferation and differentiation through multiple interacting and competing signal transduction pathways. Bone 32, 241–251 (2003).

52. De Leeuw, A. M., et al. Physiological cell bioprinting density in human bone-derived cell-laden scaffolds enhances matrix mineralization rate and stiffness under dynamic loading. Front. Bioeng. Biotechnol. 12, 1310289 (2024).

53. Rojas-Murillo, A., et al. Bioreactor-Induced Compression in Scaffolds for the Enhancement of Cartilage Regeneration: A Systematic Review. Ann Biomed Eng https://doi.org/10.1007/s10439-026-04135-4 (2026) doi:10.1007/s10439-026-04135-4.

54. Schindelin, J., et al. Fiji: an open-source platform for biological-image analysis. Nat Methods 9, 676–682 (2012).

